# Allosteric Modulation of β1 Integrin Attenuates Motor Asymmetry in the Unilateral 6-Hydroxydopamine Injury Model in Mice

**DOI:** 10.64898/2026.06.18.733264

**Authors:** Robert J. Naylor, Rehab Aljamal-Naylor

## Abstract

Parkinson’s disease (PD) is characterised by progressive dopaminergic neurodegeneration in the substantia nigra, leading to debilitating motor dysfunction. Current treatments remain largely symptomatic, highlighting the need for disease-modifying therapies. β1 integrin, implicated in neuroinflammation and trophic signalling, represents a candidate therapeutic target. We investigated whether allosteric β1 integrin modulation could attenuate motor asymmetry in the unilateral 6-hydroxydopamine (6-OHDA) mouse model of PD. Adult male C57BL/6 mice received intracerebral 6-OHDA into the substantia nigra. The anti-β1 integrin antibody JB1a (50 µg) was administered prophylactically (3 days pre-lesion) or therapeutically (3 or 7 days post-lesion). Motor asymmetry was assessed through spontaneous circling (5 min) and apomorphine-induced (0.5 mg/kg s.c.) circling (30 min). 6-OHDA induced dose-dependent contralateral circling, confirming nigrostriatal lesion. **Pre-treatment** with JB1a (3 days before 6-OHDA) reduced apomorphine-induced circling, although this did not reach statistical significance (28.5 ± 12.8, n = 4 versus 38.6 ± 7.5, n = 8; *p*>0.05). **Post-treatment at 3 days** post-lesion produced no statistically significant change in either spontaneous or apomorphine-induced circling (*p*>0.05). **Post-treatment at 7 days** post-lesion reduced apomorphine-induced circling by approximately 50%, with values returning to those of sham-operated controls (n =8-9; *p*<0.01). These findings, obtained in a murine 6-OHDA model, indicate that allosteric β1 integrin modulation attenuates lesion-induced motor asymmetry with apparent temporal specificity. As apomorphine-induced rotation reflects post-synaptic dopamine receptor supersensitivity rather than direct neuronal preservation, and as histological confirmation of dopaminergic integrity was not obtainable in this study, the present data should be interpreted as proof-of-concept behavioural evidence requiring further mechanistic and translational validation in models incorporating α-synuclein pathology. The findings are not directly generalizable to human Parkinson’s disease. The histological confirmation of lesion extent was not available and as such the behavioural findings are correspondingly interpreted as a proof-of-concept observation requiring histological replication.

## 1. Introduction

Parkinson’s disease (PD) is the second most prevalent neurodegenerative disorder globally and presents formidable therapeutic challenges (Guo et al., 2025). The disease manifests clinically through progressive motor abnormalities including tremor, bradykinesia, altered gait, muscular rigidity, and postural instability, alongside autonomic dysfunction. The principal pathological hallmark of PD is the selective and progressive loss of dopaminergic neurons in the substantia nigra pars compacta, resulting in striatal dopamine depletion that underlies the characteristic motor symptoms. The risk of Parkinsonian dementia is also recognised as the disease progresses, although the magnitude of risk remains debated (Backstrom, 2022; Leroy, 2025).

The precise causal factors for PD remain elusive; however, substantial evidence indicates that genetic susceptibility and environmental factors interact in disease pathogenesis. Mechanisms identified to date include α-synuclein aggregation (Spillantini et al., 1997), oxidative stress (Jenner, 1991), ferroptosis (Do et al., 2016), mitochondrial dysfunction (Burtscher et al., 2021), neuroinflammation (Grotemeyer et al., 2022), and gut dysbiosis (Devos et al., 2013).

Current PD therapy is predominantly symptomatic and centred on levodopa combined with carbidopa (Foltynie et al., 2024). The contemporary therapeutic pipeline, however, is increasingly focused on disease-modifying strategies, which now constitute approximately 44% of ongoing clinical trials and include α-synuclein-targeted therapies, NLRP3 inflammasome inhibitors, and GLP-1 receptor agonists (McFarthing et al., 2024).

Integrins are heterodimeric transmembrane receptors that mediate cell–ECM and cell–cell adhesion and link the ECM to intracellular cytoskeletal and signalling machinery (Luo and Springer, 2006). Their involvement in inflammatory responses and tissue remodelling has raised interest in anti-integrin reagents for neurodegenerative disorders, although direct evidence in PD is limited (Wright et al., 2007). Among integrins, β1 integrin is of particular interest: it has been identified as an alternative receptor for glial cell line-derived neurotrophic factor (GDNF), a potent dopaminergic survival factor (Cao et al., 2008), and it mediates microglial migration and morphological changes in response to neuron-released α-synuclein independently of TLR2 (Kim et al., 2014).

β1 integrin exists in multiple conformational states — bent-closed (inactive), extended-closed (primed), and extended-open (active) — that regulate ligand binding affinity and downstream signalling (Shimaoka and Springer, 2003). Allosteric modulators, unlike orthosteric blocking antibodies, bind to sites distal to the ligand-binding pocket and shift the conformational equilibrium without fully abolishing receptor function. JB1a targets the hybrid domain (amino acids 82–87) of β1 integrin, inducing a conformational change that modulates — rather than blocks — integrin activation, thereby preserving essential physiological functions while redirecting pathological signalling (Supplementary Figure S3B).

Our group has previously shown that targeting β1 integrin with allosteric modifying antibodies induces tissue repair in animal models of emphysema (Aljamal-Naylor et al., 2012) and inflammatory arthritis as well as replicative senescence (granted patent and unpublished, Al-Jamal and Harrison, 2008).

The present study tested the hypothesis that allosteric modulation of β1 integrin with JB1a, administered either before or after a unilateral nigrostriatal lesion, attenuates lesion-induced motor asymmetry in mice. We did not aim to establish dopaminergic neuroprotection per se, but to provide proof-of-concept functional evidence that may justify further mechanistic and translational investigation.

## 2. Materials and Methods

### 2.1. Animals

Adult male C57BL/6 mice (25–30 g; n= 94) were housed in groups of five (cage dimensions 266 × 425 × 185 mm; floor area 800 cm²) with free access to food (Standard Rat and Mouse No. 1) and water at 21 ± 1 °C and a 12 h light/dark cycle (lights on 07:00–19:00). All procedures were conducted under a UK Project Licence and personal licences in accordance with the UK Animals (Scientific Procedures) Act 1986. Behavioural assessments were performed between 09:30 and 18:00.

### 2.2. Randomisation and blinding

Animals were assigned to treatment groups using cage-level randomisation. Behavioural scoring was performed by a primary observer who was not formally blinded to treatment allocation (RJN); a second observer was present at the time of assessment to verify directional scoring (RAN). Formal blinding was not implemented, which we acknowledge as a limitation (Section 4.5). All assessments were video recorded.

### 2.3. Sample size

Group sizes were based on prior experience with the 6-OHDA model.

### 2.4. Surgical procedures

Surgery was performed under pentobarbitone anaesthesia (60 mg/kg, i.p.) with buprenorphine analgesia (Vetergesic®, 0.05 mg/kg, s.c.). Post-operative buprenorphine was provided in a jelly preparation in the recovery cage for voluntary intake. Anaesthetised animals were placed on a Kopf stereotaxic frame on a temperature-controlled mat with the incisor bar raised 1 mm to ensure a level skull surface from Bregma. Saline (0.5 mL, s.c., 0.9% NaCl, Baxter, UK) was administered to replenish fluid losses. The scalp was shaved and disinfected with aqueous iodine (Betadine, Seton Healthcare, UK).

Coordinates for unilateral injection into the right substantia nigra were determined from preliminary work using the atlas of Franklin and Paxinos (1997): 3.5 mm posterior to Bregma, 1.5 mm lateral to the midline, and 4.25 mm vertical from the skull surface. A burr hole was drilled, the injection delivered, and the hole sealed with dental acrylic cement. Incisions were closed with Ethicon sutures (5-0 Perma-Hand silk or coated Vicryl Plus, Johnson & Johnson). Post-operatively, enrofloxacin (Baytril®, Bayer, 0.1% v/v) was added to the drinking water and animals were monitored continuously until full recovery.

Animals receiving a second intracerebral injection were re-prepared as above, with a fresh skin incision avoiding the first site, removal of the dental cement plug, and re-injection at the same coordinates. Skin closure additionally used Dermabond (Johnson & Johnson).

All intracerebral injections used a 5 µL Hamilton syringe delivering 1 µL over 5 min, with the needle left in place for a further 5 min. Lesions were induced by 6-OHDA (Sigma H116; stabilised with ascorbic acid) dissolved in nitrogen-bubbled 0.9% saline at 1 or 2 µg/µL for the dose-effect experiment, and 2 µg/µL for the pre- and post-treatment experiments.

### 2.5. JB1a antibody

JB1a (clone JB1a, also known as J10; originally provided by J.A. Wilkins, University of Manitoba) is a mouse monoclonal antibody directed against amino acids 82–87 of the hybrid domain of human/mouse β1 integrin. The allosteric effect was determined by FRET assay as previously reported (AlJamal-Naylor et al., 2012); detailed characterisation is shown in Supplementary Figure S3C. Endotoxin testing was not performed; this is acknowledged as a limitation. The dose of 50 µg (≋ 2.5–3 mg/kg) was selected based on prior efficacy in our emphysema (AlJamal-Naylor et al., 2012) and arthritis (granted patent and unpublished) models. Intracerebral delivery was chosen to ensure direct CNS exposure, bypassing the blood–brain barrier.

### 2.6. Experimental groups and nomenclature

To clarify the experimental design, the following standardised group nomenclature is used throughout this manuscript and in all figures:

- **Sham**: surgical procedure (anaesthesia, stereotaxic placement, burr-hole drilling, dental cement sealing) without intracerebral injection.
- **Vehicle-** (1 µL or 2 µL): nitrogen-bubbled 0.9% saline injected intracerebrally at the substantia nigra coordinates, controlling for the mechanical and volumetric effects of the lesion injection.
- **Vehicle+JB1a**: diluted PBS (the JB1a reconstitution buffer) injected at the same volume and coordinates as JB1a, controlling for the antibody vehicle and the second surgical procedure.
- **6-OHDA** (1 µg or 2 µg): 6-OHDA injected intracerebrally without subsequent JB1a or treatment-vehicle administration.
- **6-OHDA + JB1a** (50 µg): 6-OHDA followed (or, in the pre-treatment cohort, preceded) by intracerebral JB1a administration.

Three independent cohorts were studied: a **pre-treatment cohort** (JB1a 3 days before 6-OHDA), a **3-day post-treatment cohort** (JB1a 3 days after 6-OHDA), and a **7-day post-treatment cohort** (JB1a 7 days after 6-OHDA). Behavioural testing was performed on day 18 (post-treatment cohorts) or day 21 (pre-treatment cohort). The full design is summarised in Supplementary Table S1.

### 2.7. Behavioural assessments

All testing was carried out in a sound-proofed, diffusely illuminated room maintained at 21 ± 1 °C. Both assessments were performed in each animal on the same testing day, with the spontaneous assessment always preceding apomorphine administration.

#### Spontaneous circling (5 min)

Animals received no acute drug treatment and were placed in an observation cage of the same dimensions as the home cage. The direction of movement was scored over a 5-min observation window by the primary observer and verified by a second observer present at the time of testing. Movement to the side opposite the hemisphere of injection was recorded as “Left” (L); movement to the same side as injection was recorded as “Right” (R). Traversing one side of the cage counted as one movement, as did a tight pivotal movement on the animal’s axis. Preliminary work confirmed that 5 min was sufficient for reliable scoring of directional bias.

#### Apomorphine-induced circling (30 min)

Following the spontaneous assessment, animals received apomorphine (0.5 mg/kg, s.c.; (S)-(+)-apomorphine hydrochloride hydrate, D043 Sigma-Aldrich; stock 0.1 mg/mL in 0.9% saline with 0.1% w/v sodium metabisulphite) and were returned to the observation cage for a 30-min observation window. Direction of movement was scored as above. Apomorphine-induced rotation in unilaterally lesioned animals reflects post-synaptic dopamine receptor supersensitivity in the denervated striatum and is therefore an indirect index of nigrostriatal asymmetry rather than a direct measure of dopaminergic neuron survival (Ungerstedt, 1971).

After the final assessment, animals were euthanised by overdose of pentobarbitone. Brains were removed and fixed in 10% w/v neutral buffered formalin for 24 h and then paraffin-embedded with the intention of subsequent immunohistochemical analysis. Unfortunately, the entire tissue archive was lost due to a technical error during processing at the Pathology Service. Consequently, no histological data are presented in this manuscript. The implications of this loss for interpretation of the behavioural findings are discussed in Section 4.5.

### 2.8. Safety and tolerability monitoring

Animals were monitored continuously during recovery and at least daily thereafter for clinical signs (posture, grooming, piloerection, seizure activity, food and water intake, and body condition). Body weight was recorded on days of all procedures according to Supplementary Table S1. There was no weight loss reported except in one animal (Supplementary Table S2). No statistically significant differences in body weight were detected between treatment groups and their matched controls at any time point (P > 0.05). Peri-operative mortality was recorded. Mortality was recorded for a total of 17 animals during the anaesthesia and was not limited to one treatment group (∼20% following double recovery surgeries). The injection site was inspected macroscopically at the time of brain removal. The full safety findings, including peri-operative outcomes, are reported in Section 3.1.

### 2.9. Statistical analysis

Sample sizes per group are reported in each figure and in Supplementary Table S2 (source data). All comparisons were between-subject (independent observations); no within-subject paired analyses were performed. Data were assessed for normality and homogeneity of variance. Group differences were analysed by one-way ANOVA with treatment as the fixed factor, with post-hoc pairwise comparisons using Tukey’s test. Where parametric assumptions were not met, the Kruskal–Wallis test with Dunn’s post-hoc was used. The three cohorts (pre-treatment, +3 d, +7 d) were analysed separately, as each tested an independent hypothesis; no global correction across cohorts was applied. Data are presented as mean ± SEM. *P* values are reported to three decimal places where available. *P* < 0.05 was considered statistically significant. Analyses were performed using GraphPad Prism 4 (Version 4.03, 2005). All the statistical analyses are provided in Supplementary Table S3.

## 3. Results

### 3.1. Safety and tolerability

Peri-operative mortality (n = 17, approximately 20%, including animals terminated on welfare grounds) occurred across cohorts and was not specific to any treatment group. Deaths occurred during or shortly after the procedure and were predominantly attributed to anaesthesia associated with double recovery surgery, consistent with expectations for this model; none were specifically attributable to JB1a. Animals showed no overt clinical signs of toxicity attributable to JB1a (no seizures, abnormal posture, piloerection, or altered grooming). Body weights included in the Supplementary Table S2). Macroscopic inspection of the injection site at brain removal showed no evidence of haemorrhage or abscess formation beyond the expected mechanical injection tract. As noted in Section 2.7, histological evaluation of the injection site was precluded by loss of the paraffin-embedded tissue archive.

### 3.2. Dose- and volume-dependence of 6-OHDA-induced motor asymmetry

#### Spontaneous circling

Spontaneous movements following unilateral intracerebral injection of Vehicle at different volumes did not differ significantly between groups (Figure 1A; *P>0.05*). Spontaneous circling did not differ significantly between 6-OHDA doses, although a trend toward greater contralateral bias at the higher dose was observed (Figure 1B; *P=0.45*).

**Figure 1.**
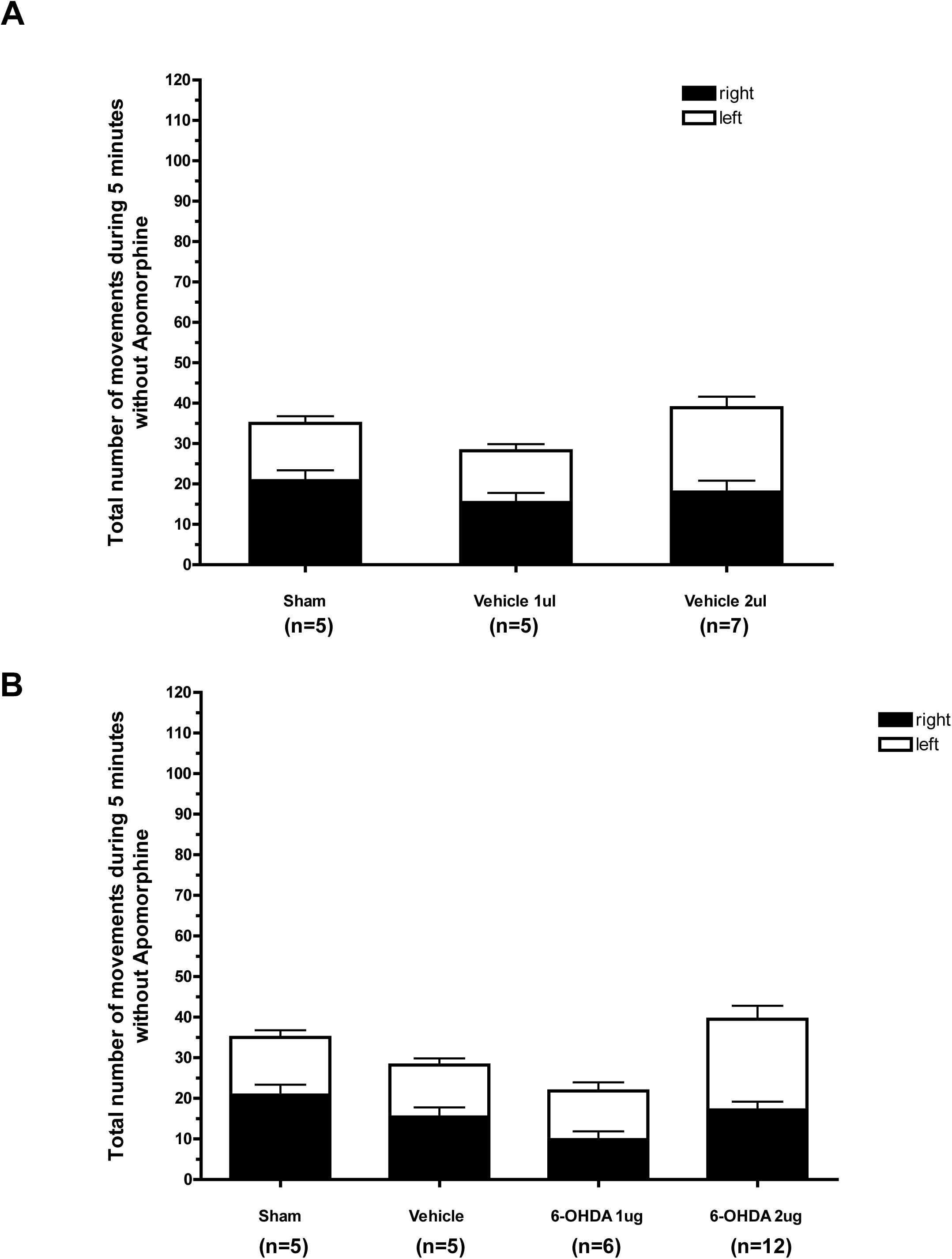
Volume- and dose-dependence of 6-OHDA on spontaneous circling. (A) Spontaneous circling assessed over a 5-min observation period 21 days after intracerebral injection of Vehicle at the indicated volumes. (B) Spontaneous circling 21 days after intracerebral 6-OHDA at the indicated doses. Data are mean + SEM; n = 5-7 per group. Differences between groups were not statistically significant (one-way ANOVA).

#### Apomorphine-induced circling

Following apomorphine administration (0.5 mg/kg s.c.), circling in Vehicle-injected animals differed significantly with injection volume (Figure 2A, *P<0.01*). 6-OHDA induced dose-dependent contralateral circling, consistent with the development of post-synaptic dopamine receptor supersensitivity in the denervated striatum (Figure 2B; *P* <0.05). The 2-µg 6-OHDA dose was used in all subsequent experiments.

**Figure 2.**
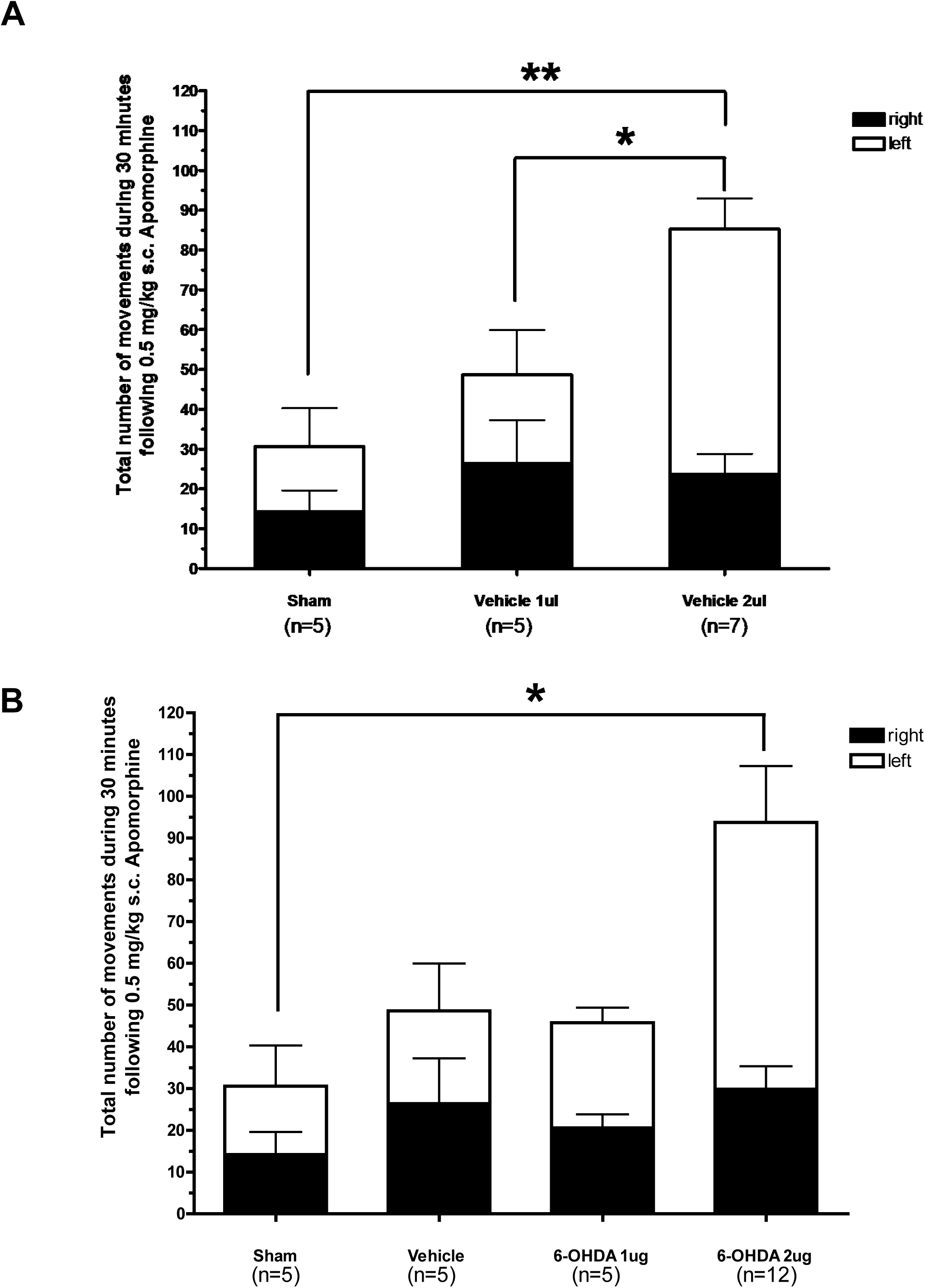
Volume- and dose-dependence of 6-OHDA on apomorphine-induced circling. (A) Apomorphine-induced (0.5 mg/kg s.c.) circling assessed over a 30-min observation period 21 days after intracerebral injection of Vehicle at the indicated volumes. (B) Apomorphine-induced circling 21 days after intracerebral 6-OHDA at the indicated doses. Data are mean + SEM; n = 5-7 per group. \**P* < 0.05 versus Vehicle (one-way ANOVA with Tukey’s post-hoc).

### 3.3. Effect of JB1a pre-treatment on motor asymmetry

In the pre-treatment cohort, JB1a (50 µg) was administered intracerebrally 3 days before 6-OHDA (2 µg). Spontaneous circling did not differ significantly between Sham, Vehicle-JB1a, 6-OHDA-only, and 6-OHDA + JB1a groups (Figure 3A; *P* >0.05; n = 4-8 per group). Apomorphine-induced circling was attenuated in 6-OHDA + JB1a animals compared with 6-OHDA-only animals but failed to achieve statistical significance (Figure 4A).

**Figure 3.**
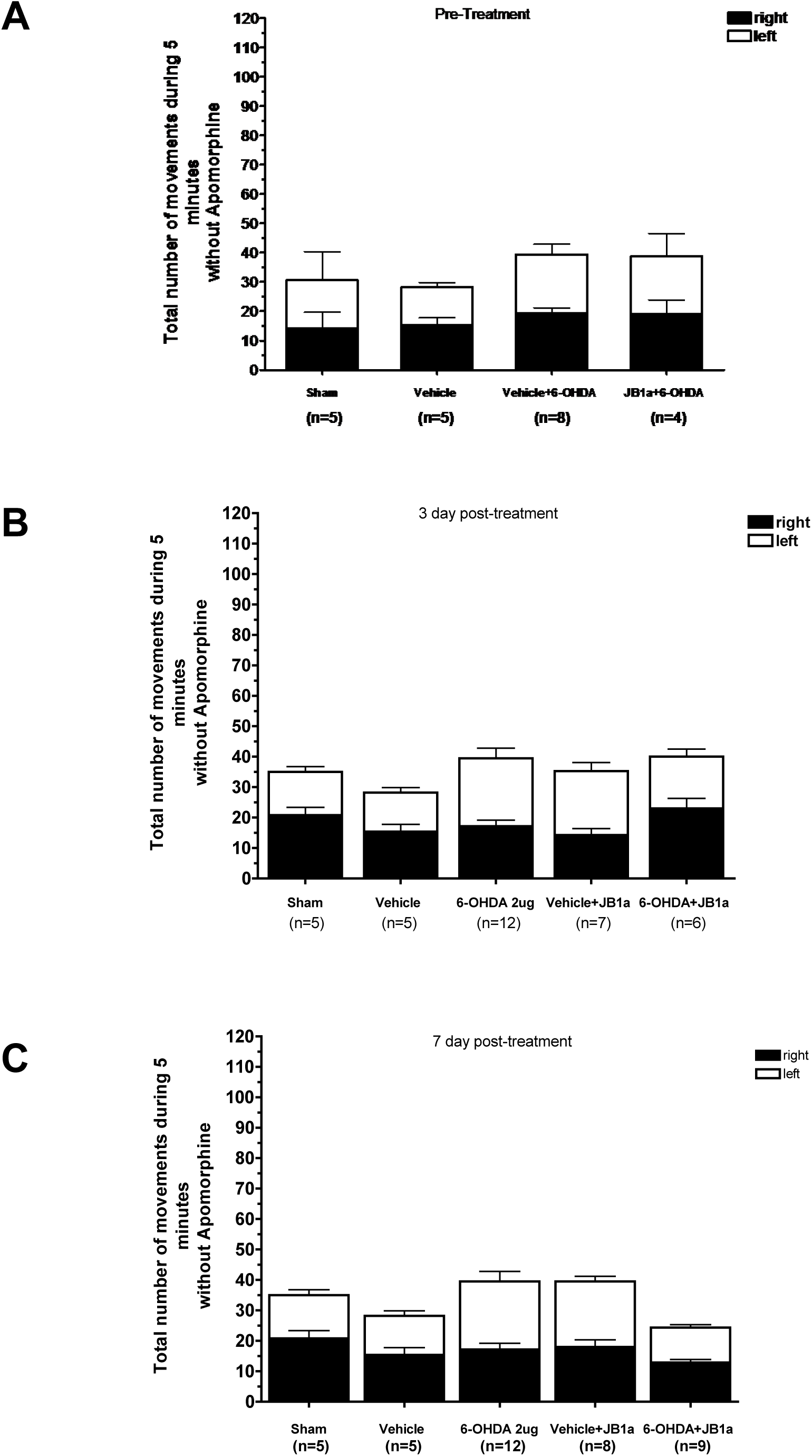
Effect of JB1a on spontaneous circling. Spontaneous circling (5-min observation) in the (A) pre-treatment, (B) 3-day post-treatment, and (C) 7-day post-treatment cohorts. Groups: Sham, Vehicle, 6-OHDA (2 µg), Vehicle + JB1a, 6-OHDA + JB1a (50 µg). Data are mean + SEM; n = 4-12 per group. \**P* < 0.05 versus 6-OHDA (one-way ANOVA with Tukey’s post-hoc).

**Figure 4.**
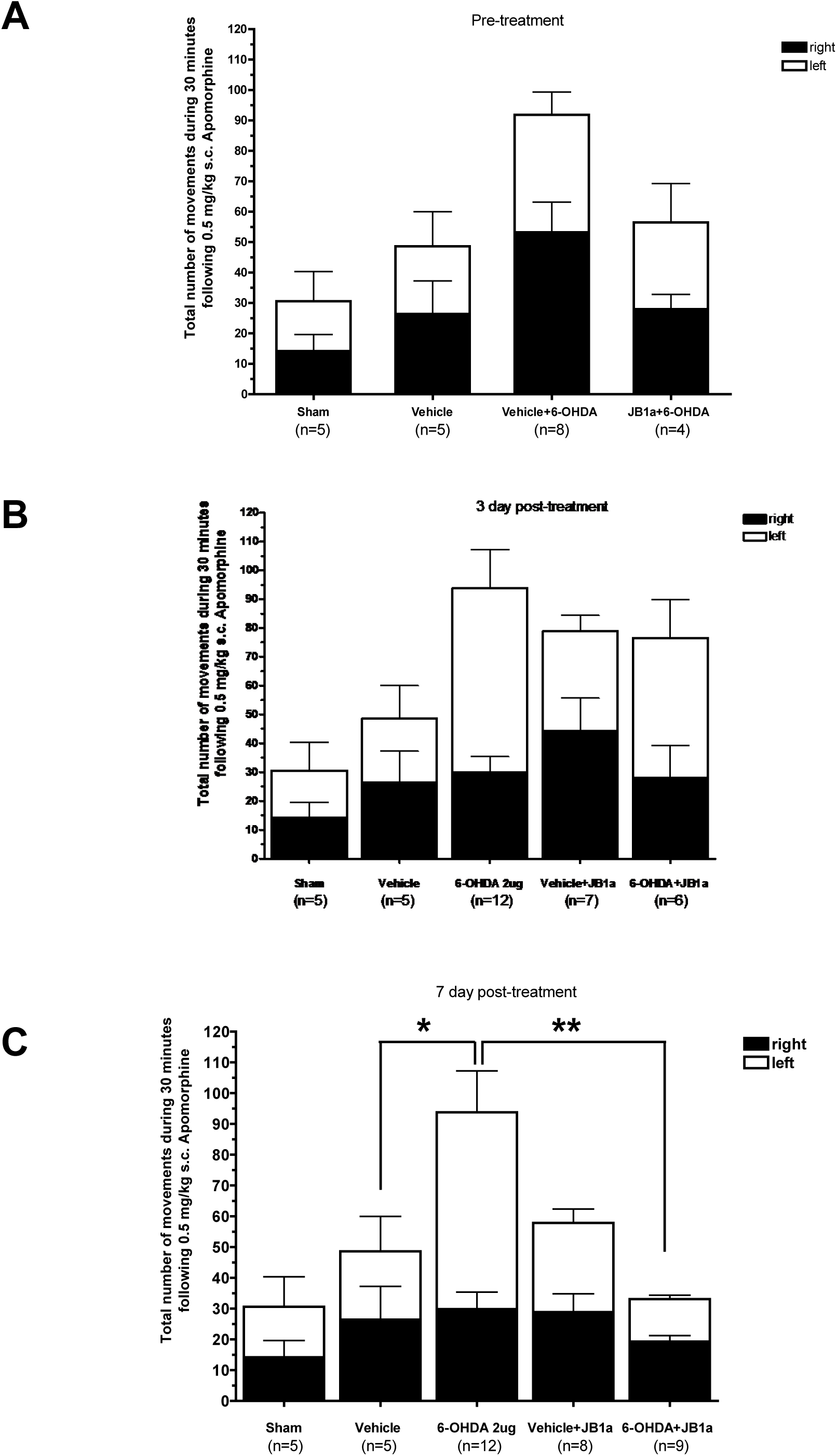
Effect of JB1a on apomorphine-induced circling. Apomorphine-induced (0.5 mg/kg s.c.) circling assessed over a 30-min observation period in the (A) pre-treatment, (B) 3-day post-treatment, and (C) 7-day post-treatment cohorts. Groups: Sham, Vehicle, 6-OHDA (2 µg), Vehicle + JB1a, 6-OHDA + JB1a (50 µg). Data are mean + SEM; n = 4-12 per group. \**P* < 0.05 versus 6-OHDA (one-way ANOVA with Tukey’s post-hoc).

### 3.4. Effect of JB1a post-treatment at 3 days on motor asymmetry

JB1a administered 3 days after 6-OHDA produced no statistically significant change in spontaneous circling (Figure 3B; *P>0.05*) apomorphine-induced circling (Figure 4B; *P* >0.05) compared with the 6-OHDA-only group.

### 3.5. Effect of JB1a post-treatment at 7 days on motor asymmetry

In contrast to the 3-day post-treatment cohort, JB1a administered 7 days after 6-OHDA significantly reduced both spontaneous circling (Figure 3C; approximately 50% reduction in contralateral bias; *P<0.05*; n = 5-12) and apomorphine-induced circling (Figure 4C; approximately 50% reduction; *P* = 0.003). In both assessments, values in the 6-OHDA + JB1a group returned toward those of the Sham group.

## 4. Discussion

### 4.1. Summary of findings

In this proof-of-concept study, allosteric modulation of β1 integrin with the JB1a antibody attenuated lesion-induced motor asymmetry in the unilateral 6-OHDA mouse model of PD. Two findings are noteworthy. First, pre-treatment with JB1a 3 days before 6-OHDA showed a trend towards reducing apomorphine-induced rotation. Second, post-treatment with JB1a was effective at 7 days, but not at 3 days, after the lesion. The dose-dependent induction of contralateral circling by 6-OHDA confirms successful unilateral nigrostriatal lesioning, consistent with established mechanisms of dopamine receptor supersensitivity (Ungerstedt, 1971).

### 4.2. Interpretation of behavioural outcomes

We interpret the present findings cautiously. Apomorphine-induced rotation in unilaterally lesioned animals reflects post-synaptic D1/D2 receptor supersensitivity in the denervated striatum and is an *indirect* index of nigrostriatal asymmetry rather than a direct measure of dopaminergic neuron preservation. The observed reduction in apomorphine-induced circling following JB1a may therefore reflect (i) preservation of pre-synaptic dopaminergic input, thereby reducing the magnitude of supersensitivity that develops; (ii) modulation of post-synaptic receptor adaptations; or (iii) a combination of both mechanisms. The present data cannot distinguish between these possibilities. We have correspondingly avoided the language of “neuroprotection” and “reversal” in describing our findings, preferring “attenuation of motor asymmetry”.

### 4.3. Mechanistic considerations

The mechanistic basis for β1 integrin involvement in PD-relevant pathology is supported by emerging evidence linking this receptor to multiple disease-relevant processes. β1 integrin mediates microglial migration and morphological responses to neuron-released α-synuclein independently of TLR2 (Kim et al., 2014), positioning it within the neuroinflammatory cascade increasingly recognised as central to PD progression (Grotemeyer et al., 2022). β1 integrin has also been identified as an alternative receptor for GDNF (Cao et al., 2008), a potent dopaminergic survival factor with a long but troubled history of clinical translation in PD (Gill et al., 2003). Either or both of these mechanisms could contribute to the behavioural effects observed here, but direct mechanistic confirmation in the present model was not undertaken.

### 4.4. Temporal pattern of post-treatment efficacy

The lack of effect at 3 days post-lesion, contrasted with significant attenuation at 7 days, may reflect the temporal evolution of secondary injury after 6-OHDA. The acute phase of excitotoxicity and oxidative stress is followed by a subacute inflammatory phase characterised by sustained microglial activation (Marinova-Mutafchieva et al., 2009; Walsh et al., 2011). The 7-day window may correspond to a critical period during which β1 integrin modulation can interrupt these secondary processes. Alternatively, ongoing inflammation at 3 days could compromise the activity or local availability of the antibody (Tian et al., 2025).

These hypotheses require direct experimental testing.

### 4.5. Limitations

Several limitations of this study must be acknowledged explicitly:

- **Loss of histological tissue.** The paraffin-embedded brain tissue from this study was lost due to a technical error during processing at the Pathology Service. Consequently, no immunohistochemical quantification of TH-positive neurons in the substantia nigra or TH-positive terminal density in the striatum is available. This is the most significant limitation of the present work and precludes definitive conclusions regarding dopaminergic preservation. Replication studies with histological confirmation would be required. Therefore, the absence of TH confirmation is a primary limitation and that failed or partial lesions cannot be excluded. Therefore, the conclusion is hypothesis-generating.
- **Restricted behavioural battery.** Functional assessment was limited to spontaneous and apomorphine-induced circling, which capture lateralised motor asymmetry but do not interrogate fine motor coordination, balance, gait, skilled forelimb use, or non-motor symptoms. Future studies should incorporate complementary assays such as the rotarod, cylinder test, stepping test, beam traversal, and automated gait analysis (e.g., CatWalk, DigiGait), and should include cognitive and affective endpoints.
- **Acute neurotoxin model.** The 6-OHDA model represents an acute neurotoxic insult and does not recapitulate the progressive, multisystem pathology of human PD or its cardinal feature of α-synuclein aggregation. However, to date, it is used for early assessements of efficacy of novel targets in PD. Validation in α-synuclein-based models (e.g., preformed fibril seeding, AAV-mediated overexpression, transgenic lines) is required before stronger conclusions can be drawn.
- **Statistical and methodological reporting.** Whilst no formal power calculation was performed; group sizes were based on prior experience with the model. Blinding of the behavioural observer was not formally implemented. Body weight was systematically recorded across all cohorts to ensure animal welfare is not compromised.
- **Delivery route.** Intracerebral injection is not yet clinically viable for antibody-based chronic therapy. Brain-penetrant formulations or alternative delivery strategies (e.g., receptor-mediated transcytosis, small-molecule allosteric modulators) would be required for translational development.
- **Single-dose design.** Only a single dose of JB1a (50 µg) was tested. Dose–response characterisation is needed. However, in previous models, we identified 3 mg/kg as a suitable dose.
- **Generalisability.** All findings derive from male C57BL/6 mice and are not directly generalisable to human PD. Sex differences and age effects were not assessed.

### 4.6. Future directions

Building on this proof-of-concept work, future studies will (i) replicate the behavioural findings with complete histological characterisation (TH immunohistochemistry, stereological quantification, terminal density); (ii) test efficacy in α-synuclein-based models; (iii) characterise the dose–response relationship and therapeutic window; (iv) include both sexes and aged animals; (v) employ a comprehensive motor and non-motor behavioural battery; and (vi) develop brain-penetrant formulations or small-molecule allosteric modulators suitable for systemic administration.

## 5. Conclusions

This study provides initial proof-of-concept behavioural evidence that allosteric modulation of β1 integrin with the JB1a antibody attenuates lesion-induced motor asymmetry in the unilateral 6-OHDA mouse model of PD, with apparent temporal specificity favouring pre-treatment and delayed (7-day) post-treatment paradigms. Because the present data are based exclusively on circling behaviour, lack histological confirmation of dopaminergic integrity, and derive from an acute neurotoxin model in male mice, they are not directly generalisable to human PD and cannot be interpreted as definitive evidence of neuroprotection or disease modification. The findings nevertheless justify further mechanistic investigation, validation in α-synuclein-based models, and the development of clinically viable delivery strategies. While no overt adverse effects were observed in this study, the safety profile of β1 integrin modulation in the CNS will require comprehensive toxicological evaluation before any clinical translation.

## Abbreviations

6-OHDA: 6-hydroxydopamine
ANOVA: analysis of variance
CNS: central nervous system
ECM: extracellular matrix
FAK: focal adhesion kinase
GDNF: glial cell line-derived neurotrophic factor
IHC: immunohistochemistry
JB1a: monoclonal antibody clone JB1a (also J10)
MAPK: mitogen-activated protein kinase
PD: Parkinson’s disease
PBS: phosphate-buffered saline
SEM: standard error of the mean
TH: tyrosine hydroxylase
TLR2: Toll-like receptor 2
FRET: fluorescence resonance energy transfer.

## Conflict of Interest Statement

R. Aljamal-Naylor and the late Robert J. Naylor are shareholders of Avipero Ltd. and Avipero Bio, which are developing anti-β1 integrin therapies. R. Aljamal-Naylor and D.J. Harrison are co-inventors of intellectual property relating to the therapeutic effects of β1 integrin modulation (WO2005037313, 17 October 2003; WO2008104808, 27 February 2007), owned by R. Aljamal-Naylor and Robert J. Naylor. The funders had no role in study design, data collection and analysis, decision to publish, or preparation of the manuscript. This conflict of interest is mirrored in the cover letter accompanying this submission.

## Author Contributions

R. AlJamal-Naylor and the late Robert J. Naylor contributed equally to the intellectual property, scientific input, study design, conduct of experiments. R. Aljamal-Naylor was solely responsible for the manuscript preparation.

## Funding

This work was supported in part by the Chief Scientist Office, Scottish Government (CZB4/602), and by Avipero Ltd.

## Acknowledgements

We thank Professor J.A. Wilkins (University of Manitoba) for providing the JB1a antibody. We acknowledge with regret the loss of paraffin-embedded brain tissue at the Pathology Service/Edinburgh University, which precluded planned histological analyses. ChatGPT and ClaudeAI were used solely for language editing of the Abstract, Introduction, and Discussion sections; it was not used for scientific input, data analysis, or interpretation.

## Data Availability

Individual animal-level data underlying all figures are provided as Supplementary Table S2. Additional materials are available from the corresponding author upon reasonable request.

## Supplementary Materials

**Supplementary Table S1.**
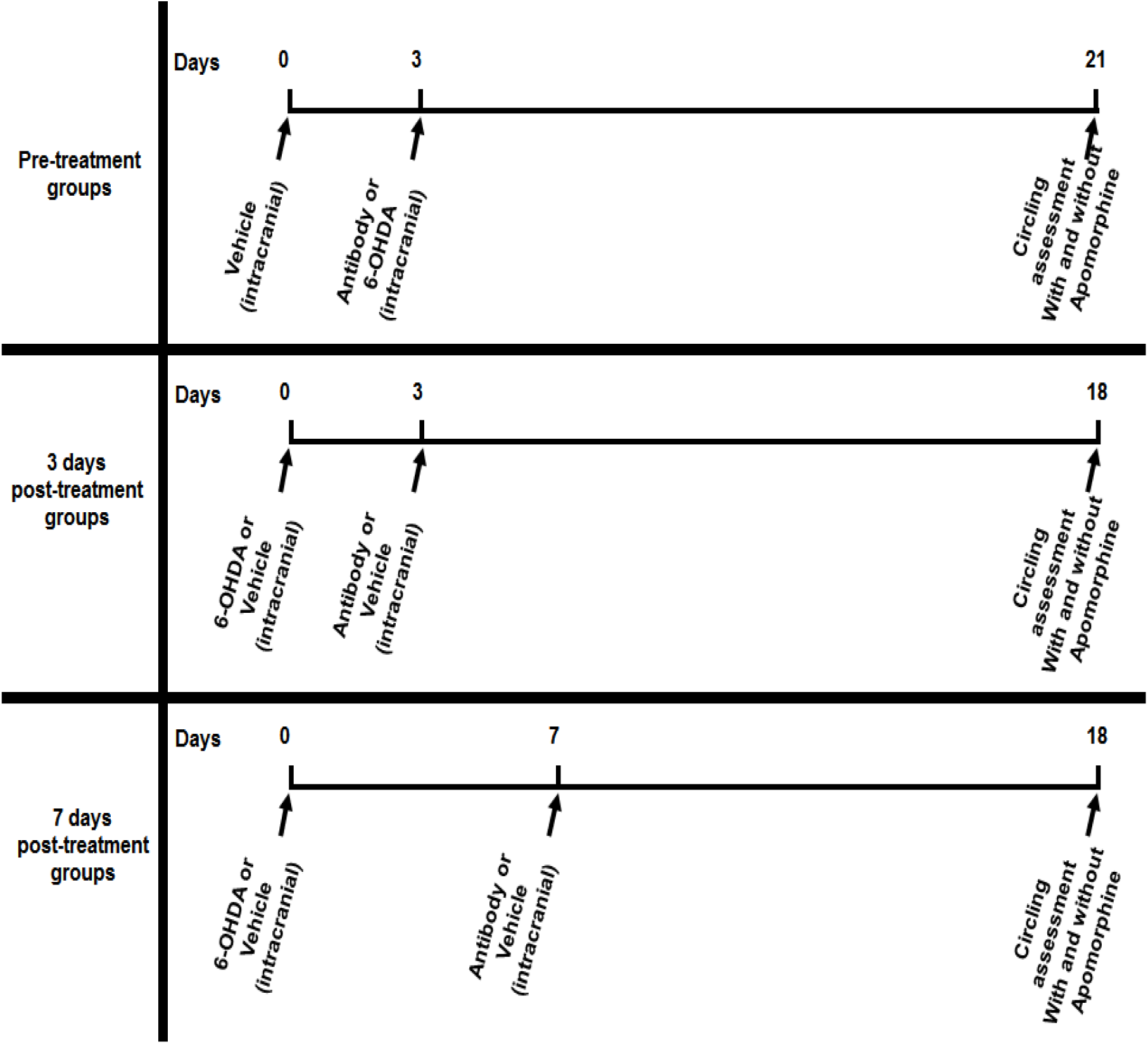
Experimental design. Schematic representation of the three independent cohorts. Pre-treatment cohort: animals received intracerebral JB1a (50 µg) or Vehicle on day −3 and 6-OHDA (2 µg) or Vehicle on day 0; behavioural testing on day 21. 3-day post-treatment cohort: 6-OHDA or Vehicle on day 0; JB1a or Vehicle on day 3; testing on day 18. 7-day post-treatment cohort: 6-OHDA or Vehicle on day 0; JB1a or Vehicle on day 7; testing on day 18. All injections were intracerebral into the right substantia nigra. Both spontaneous (5 min) and apomorphine-induced (0.5 mg/kg s.c., 30 min) circling were assessed on the same testing day, with the spontaneous assessment preceding apomorphine.

**Supplementary Table S2.** Individual animal-level source data for all figure panels, including means, SD, SEMs, n=number of animals in each group.

| Animal number | No drug administration:<br>Behavioural observation only<br>(5 minute period) |  | Asymmetry<br>(30 minute period) | Apomorphine administration<br>“Circling (directional) behaviour” (30 minute) |  | Weight before<br>(g) | Weight after<br>(g) |
| --- | --- | --- | --- | --- | --- | --- | --- |
|  | Right movements | Left movements |  | Right movement | Left movement |  |  |
| 1 | 26 | 11 |  | 17 | 50 |  |  |
| 2 | 26 | 12 |  | 34 | 27 | 27 | 27 |
| 3 | 17 | 19 |  | 9 | 3 | 25 | 25 |
| 4 | 22 | 18 |  | 3 | 1 | 25 | 25 |
| 5 | 13 | 11 |  | 8 | 1 | 24 | 24 |
| mean | 20.8 | 14.2 |  | 14.2 | 16.4 | 25.25 | 25.25 |
| SD | 5.7183914 | 3.9623226 |  | 12.153189 | 21.766947 | 1.2583057 | 1.2583057 |
| SEM | 2.5573424 | 1.7720045 |  | 5.4350713 | 9.7344748 | 0.6291529 | 0.6291529 |

Vehicle (1µl)
| Animal number | No drug administration:<br>Behavioural observation only<br>(5 minute period) |  | Asymmetry<br>(30 minute period) | Apomorphine administration<br>“Circling (directional) behaviour” (30 minute) |  | Weight before<br>(g) | Weight after<br>(g) |
| --- | --- | --- | --- | --- | --- | --- | --- |
|  | Right movements | Left movements |  | Right movement | Left movement |  |  |
| 1 | 16 | 10 |  | 4 | 6 | 30 | 25 |
| 2 | 23 | 12 |  | 62 | 5 | 26 | 25 |
| 3 | 12 | 9 |  | 7 | 66 | 25 | 25 |
| 4 | 17 | 18 |  | 20 | 22 | 28 | 26 |
| 5 | 9 | 15 |  | 39 | 12 | 25 | 26 |
| mean | 15.4 | 12.8 |  | 26.4 | 22.2 | 26.8 | 25.4 |
| SD | 5.3197744 | 3.7013511 |  | 24.213633 | 25.400787 | 2.1679483 | 0.5477226 |
| SEM | 2.3790755 | 1.6552945 |  | 10.828666 | 11.359577 | 0.969536 | 0.244949 |

Vehicle (2µl)
| Animal number | No drug administration:<br>Behavioural observation only<br>(5 minute period) |  | Asymmetry<br>(30 minute period) | Apomorphine administration<br>“Circling (directional) behaviour” (30 minute) |  | Weight before<br>(g) | Weight after<br>(g) |
| --- | --- | --- | --- | --- | --- | --- | --- |
|  | Right movements | Left movements |  | Right movement | Left movement |  |  |
| 1 | 24 | 31 |  | 15 | 75 | 23 | 25 |
| 2 | 9 | 28 |  | 12 | 41 | 27 | 26 |
| 3 | 17 | 15 |  | 13 | 89 | 24 | 24 |
| 4 | 27 | 23 |  | 22 | 45 | 24 | 28 |
| 5 (1) | 14 | 23 |  | 28 | 60 | 24 | 25 |
| 6 (2) | 10 | 13 |  | 25 | 40 | 24 | 24 |
| 7 (3) | 25 | 13 |  | 51 | 81 | 20 | 21 |
| mean | 18 | 20.857143 |  | 23.714286 | 61.571429 | 23.714286 | 24.714286 |
| SD | 7.393691 | 7.3127417 |  | 13.5119 | 20.313027 | 2.0586635 | 2.1380899 |
| SEM | 2.7945525 | 2.7639565 |  | 5.107018 | 7.6776024 | 0.7781016 | 0.808122 |

6-OHDA (1µg)
| Animal number | No drug administration:<br>Behavioural observation only<br>(5 minute period) |  | Asymmetry<br>(30 minute period) | Apomorphine administration<br>“Circling (directional) behaviour” (30 minute) |  | Weight before<br>(g) | Weight after<br>(g) |
| --- | --- | --- | --- | --- | --- | --- | --- |
|  | Right movements | Left movements |  | Right movement | Left movement |  |  |
| 1 | 12 | 13 |  | 26 | 31 | 26 | 28 |
| 2 | 17 | 15 |  | Died inj | Died inj | 29 | 29 |
| 3 | 7 | 18 |  | 28 | 18 | 26 | 25 |
| 4 | 4 | 3 |  | 17 | 18 | 26 | 26 |
| 5 | 6 | 10 |  | 10 | 36 | 27 | 28 |
| 6 (1) | 13 | 13 |  | 22 | 23 | 26 | 25 |
| mean | 9.8333333 | 12 |  | 20.6 | 25.2 | 26.666667 | 26.833333 |
| SD | 4.9564772 | 5.138093 |  | 7.2663608 | 8.043631 | 1.2110601 | 1.7224014 |
| SEM | 2.0234734 | 2.0976177 |  | 3.2496154 | 3.5972211 | 0.4944132 | 0.7031674 |

6-OHDA (2µg)
| Animal number | No drug administration:<br>Behavioural observation only<br>(5 minute period) |  | Asymmetry<br>(30 minute period) | Apomorphine administration<br>“Circling (directional) behaviour” (30 minute) |  | Weight before (g) | Weight after (g) |
| --- | --- | --- | --- | --- | --- | --- | --- |
|  | Right movements | Left movements |  | Right movement | Left movement |  |  |
| 1 | 23 | 14 |  | 37 | 45 |  | 21 |
| 2 | 16 | 33 |  | 35 | 70 |  | 21 |
| 3 | 16 | 21 |  | 27 | 67 |  | 22 |
| 4 | 18 | 27 |  | 54 | 130 |  | 22 |
| 5 | 17 | 16 |  | 9 | 41 |  | 20 |
| 6 (1) | 22 | 36 |  | 57 | 59 |  | 27 |
|  | 22 | 23 |  | 21 | 42 | 27 | 28 |
|  | 32 | 42 |  | 62 | 30 | 26 | 26 |
|  | 11 | 9 |  | 19 | 86 | 26 | 28 |
|  | 5 | 2 |  | 16 | 11 | 26 | 26 |
|  | 12 | 25 |  | 9 | 171 | 28 | 27 |
|  | 12 | 20 |  | 12 | 15 | 27 | 28 |
| mean | 17.166667 | 22.333333 |  | 29.833333 | 63.916667 | 26.666667 | 24.666667 |
| SD | 7.0302377 | 11.380472 |  | 19.106559 | 46.806096 | 0.8164966 | 3.1718458 |
| SEM | 2.0294548 | 3.2852594 |  | 5.5155886 | 13.511756 | 0.3333333 | 0.915633 |

| Animal number | No drug administration:<br>Behavioural observation only<br>(5 minute period) |  | Asymmetry<br>(30 minute period) | Apomorphine administration<br>“Circling (directional) behaviour” (30 minute) |  | Weight before (g) | Weight after (g) |
| --- | --- | --- | --- | --- | --- | --- | --- |
|  | Right movements | Left movements |  | Right movement | Left movement |  |  |
| 1 | 19 | 17 |  | 34 | 46 | 23 | 22 |
| 2 | 7 | 20 |  | 110 | 19 | 24 | 20 |
| 3 | 20 | 16 |  | 27 | 51 | 23 | 21 |
| 4 | 15 | 30 |  | 46 | 11 | 23 | 21 |
| 5 | 10 | 33 |  | 43 | 36 | 21 | 20 |
| 6 (1) | 9 | 16 |  | 17 | 35 | 22 | 20 |
| 7 (2) | 20 | 15 |  | 32 | 45 | 19 | 19 |
| mean | 14.285714 | 21 |  | 44.142857 | 34.714286 | 22.142857 | 20.428571 |
| SD | 5.589105 | 7.393691 |  | 30.612789 | 14.772884 | 1.6761634 | 0.9759001 |
| SEM | 2.1124831 | 2.7945525 |  | 11.570547 | 5.5836252 | 0.6335302 | 0.3688556 |

| Animal number | No drug administration:<br>Behavioural observation only<br>(5 minute period) |  | Asymmetry<br>(30 minute period) | Apomorphine administration<br>“Circling (directional) behaviour” (30 minute) |  | Weight before (g) | Weight after (g) |
| --- | --- | --- | --- | --- | --- | --- | --- |
|  | Right movements | Left movements |  | Right movement | Left movement |  |  |
| 1 | 22 | 12 |  | 12 | 68 | 29 | 23 |
| 2 | 35 | 23 |  | 6 | 89 | 24 | 23 |
| 3 | 11 | 10 |  | 20 | 11 | 23 | 22 |
| 4 | 23 | 13 |  | 27 | 35 | 26 | 24 |
| 5 | 28 | 24 |  | 21 | 15 | 26 | 23 |
| 6 | 19 | 20 |  | 82 | 73 | 29 | 27 |
| mean | 23 | 17 |  | 28 | 48.5 | 26.166667 | 23.666667 |
| SD | 8.1240384 | 6.0663004 |  | 27.45906 | 32.654249 | 2.4832774 | 1.7511901 |
| SEM | 3.3166248 | 2.4765567 |  | 11.210114 | 13.331041 | 1.0137938 | 0.7149204 |

S2. Vehicle+ JB1a (7day)
| Animal number | No drug administration: Behavioural observation only (5 minute period) |  | Asymmetry (30 minute period) | Apomorphine administration "Circling (directional) behaviour" (30 minute) |  | Weight before (g) | Weight after (g) |
| --- | --- | --- | --- | --- | --- | --- | --- |
|  | Right movements | Left movements |  | Right movement | Left movement |  |  |
| 1 | 20 | 16 |  | 18 | 31 | 27 | 27 |
| 2 | 28 | 22 |  | 51 | 105 | 30 | 28 |
| 3 | 23 | 21 |  | 23 | 32 | 28 | 28 |
| 4 | 11 | 23 |  | 57 | 16 | 26 | 25 |
| 5 (1) | 13 | 15 |  | 14 | 19 | 28 | 28 |
| 6 (2) | 24 | 31 |  | 11 | 41 | 32 | 31 |
| 7 (3) | 11 | 23 |  | 31 | 48 | 27 | 27 |
| 8 (4) | 14 | 21 |  | 26 | 16 | 33 | 33 |
| mean | 18 | 21.5 |  | 28.875 | 38.5 | 28.875 | 28.375 |
| SD | 6.5900358 | 4.8989795 |  | 16.847531 | 29.30139 | 2.5319388 | 2.5035689 |
| SEM | 2.3299295 | 1.7320508 |  | 5.9565015 | 10.359606 | 0.8951756 | 0.8851453 |

6-OHDA+ JB1a (7day)
| Animal number | No drug administration: Behavioural observation only (5 minute period) |  | Asymmetry (30 minute period) | Apomorphine administration "Circling (directional) behaviour" (30 minute) |  | Weight before (g) | Weight after (g) |
| --- | --- | --- | --- | --- | --- | --- | --- |
|  | Right movements | Left movements |  | Right movement | Left movement |  |  |
| 1 | 12 | 13 |  | 13 | 15 | 28 | 29 |
| 2 | 14 | 14 |  | 24 | 20 | 28 | 24 |
| 3 | 9 | 8 |  | 14 | 13 | 27 | 23 |
| 4 | 16 | 13 |  | 23 | 9 | 25 | 25 |
| 5 (1) | 10 | 7 |  | 12 | 17 | 24 | 28 |
| 6 (2) | 10 | 10 |  | 28 | 13 | 26 | 28 |
| 7 (3) | 15 | 10 |  | 22 | 16 | 24 | 26 |
| 8 (4) | 17 | 12 |  | 15 | 12 | 30 | 28 |
| 9 (5) | 13 | 16 |  | 23 | 9 | 27 | 28 |
| mean | 12.888889 | 11.444444 |  | 19.333333 | 13.777778 | 26.555556 | 26.555556 |
| SD | 2.8480012 | 2.9202359 |  | 5.8309519 | 3.6324158 | 2.0069324 | 2.1278576 |
| SEM | 0.9493337 | 0.973412 |  | 1.9436506 | 1.2108053 | 0.6689775 | 0.7092859 |

JB1a+ Vehicle (3day)
| Animal number | No drug administration: Behavioural observation only (5 minute period) |  | Asymmetry (30 minute period) | Apomorphine administration "Circling (directional) behaviour" (30 minute) |  | Weight before (g) | Weight after (g) |
| --- | --- | --- | --- | --- | --- | --- | --- |
|  | Right movements | Left movements |  | Right movement | Left movement |  |  |
| 1 | 28 | 43 |  | 25 | 62 | 26 | 26.8 |
| 2 | 11 | 10 |  | 17 | 34 | 27 | 34.3 |
| 9 | 10 | 11 |  | 40 | 4 | 27.4 | 26.1 |
| 10 | 27 | 15 |  | 30 | 14 | 26.4 | 26.3 |
| mean | 19 | 19.75 |  | 28 | 28.5 | 26.7 | 28.375 |
| SD | 9.8319208 | 15.649814 |  | 9.6263527 | 25.57994 | 0.6218253 | 3.9609553 |
| SEM | 4.9159604 | 7.8249068 |  | 4.8131764 | 12.78997 | 0.3109126 | 1.9804776 |

JB1a+ 6-OHDA (3day)
| Animal number | No drug administration: Behavioural observation only (5 minute period) |  | Asymmetry (30 minute period) | Apomorphine administration "Circling (directional) behaviour" (30 minute) |  | Weight before (g) | Weight after (g) |
| --- | --- | --- | --- | --- | --- | --- | --- |
|  | Right movements | Left movements |  | Right movement | Left movement |  |  |
| S3 | 13 | 23 |  | 93 | 88 | 29 | 27.4 |
| S4 | 24 | 18 |  | 96 | 40 | 28 | 29 |
| S5 | 28 | 16 |  | 46 | 28 | 31.7 | 32.2 |
| S6 | 16 | 9 |  | 50 | 22 | 25 | 24 |
| S7 | 24 | 43 |  | 56 | 28 | 26.5 | 26 |
| S8 | 14 | 15 |  | 24 | 38 | 28 | 26.5 |
| S9 | 16 | 19 |  | 23 | 40 | 25 | 24.6 |
| S10 | 19 | 18 |  | 38 | 25 | 30.7 | 30.4 |
| mean | 19.25 | 20.125 |  | 53.25 | 38.625 | 27.9875 | 27.5125 |
| SD | 5.4707012 | 10.063193 |  | 27.983413 | 21.14533 | 2.4561512 | 2.8452655 |
| SEM | 1.934185 | 3.5578761 |  | 9.8936307 | 7.4760033 | 0.8683806 | 1.0059533 |

**Supplementary Table S3.**
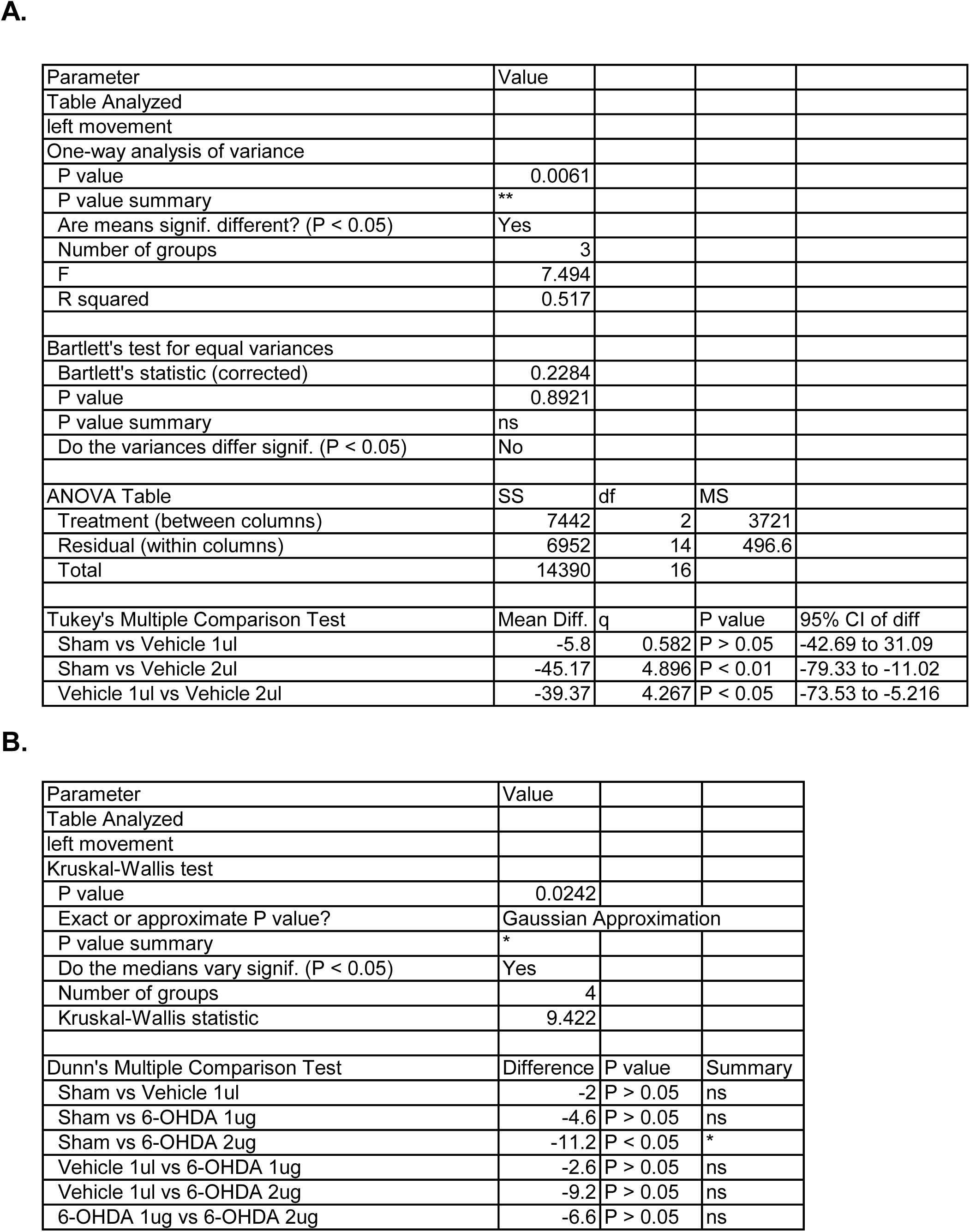
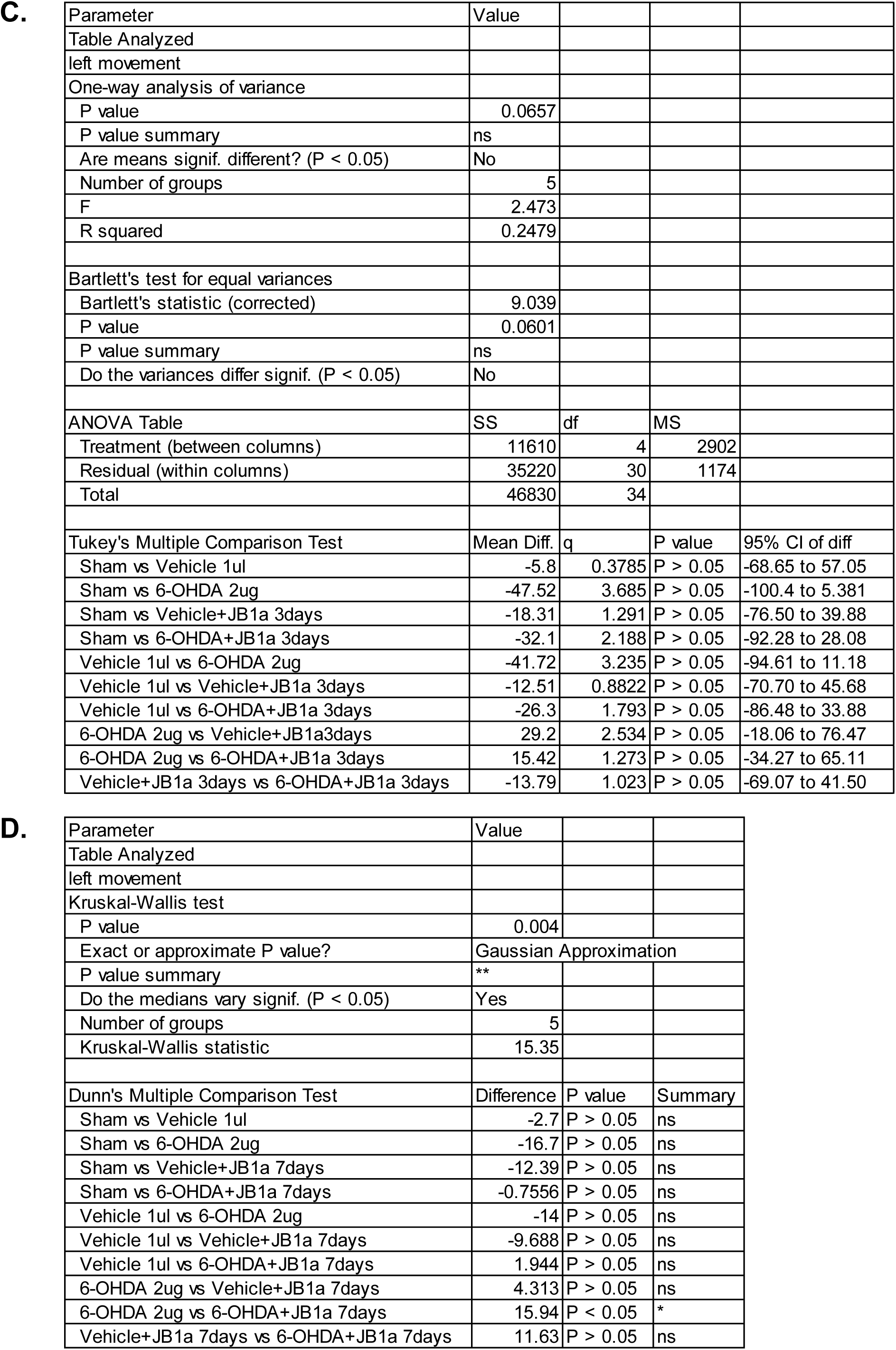
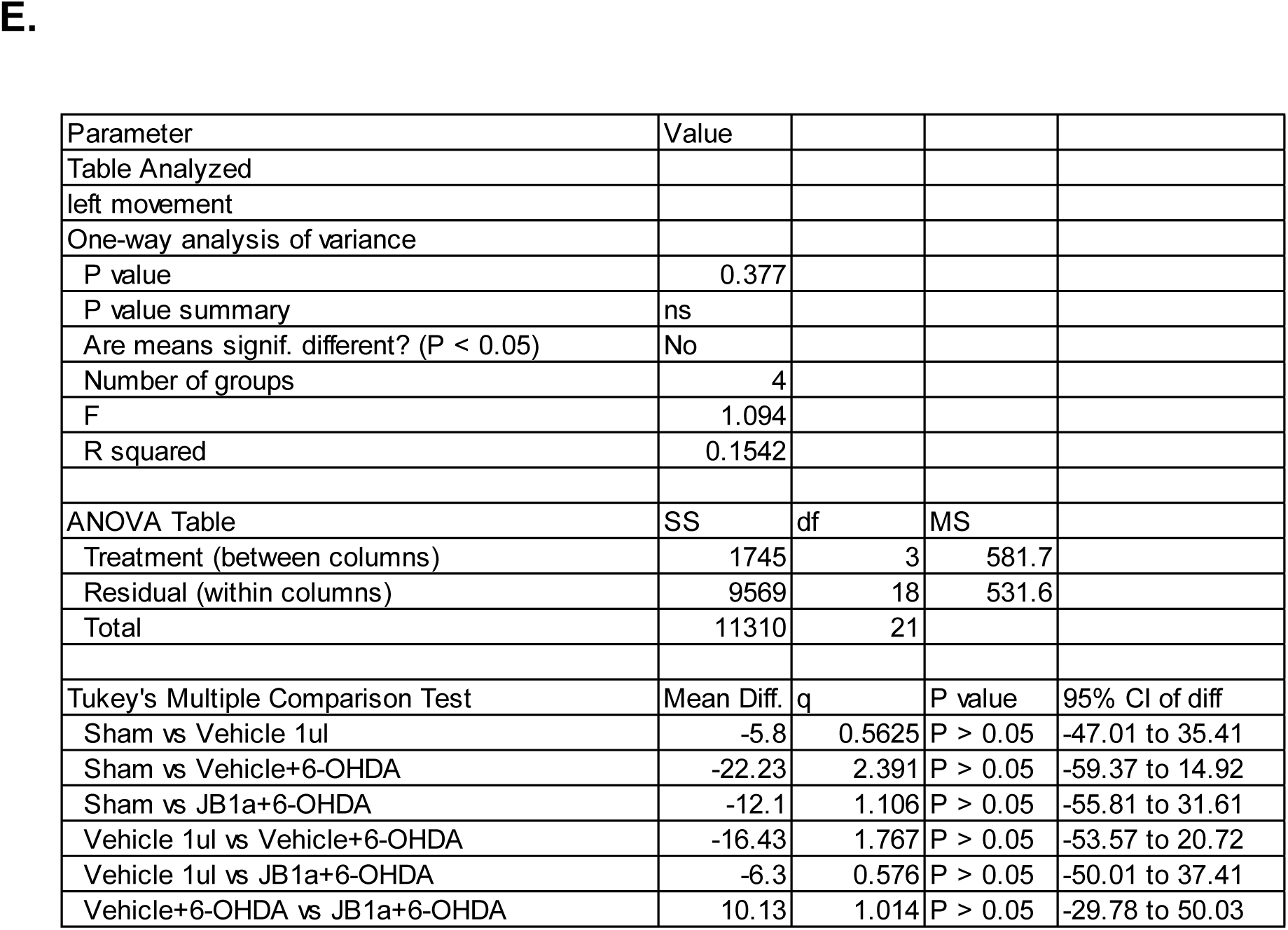
Statistical analyses for all the cohorts as described in Supplementary Table S1. (A) Volume-effect on apomorphine-induced left circling (0.5 mg/kg s.c.) assessed over a 30-min observation period 21 days after intracerebral injection of Vehicle at the indicated volumes. (B) 6-OHDA-effect on apomorphine-induced left circling (0.5 mg/kg s.c.) assessed over a 30-min observation period 21 days after intracerebral injection of 6-OHDA at the indicated doses. Apomorphine-induced (0.5 mg/kg s.c.) circling assessed over a 30-min observation period in the (C) 3-day post-treatment, (D) 7-day post-treatment and (E) pre-treatment cohorts.

A.
| Parameter | Value |  |  |  |
| --- | --- | --- | --- | --- |
| Table Analyzed |  |  |  |  |
| left movement |  |  |  |  |
| One-way analysis of variance |  |  |  |  |
| P value | 0.0061 |  |  |  |
| P value summary | ** |  |  |  |
| Are means signif. different? ($P < 0.05$) | Yes | | | |
| Number of groups | 3 |  |  |  |
| F | 7.494 |  |  |  |
| R squared | 0.517 |  |  |  |
| Bartlett's test for equal variances |  |  |  |  |
| Bartlett's statistic (corrected) | 0.2284 |  |  |  |
| P value | 0.8921 |  |  |  |
| P value summary | ns |  |  |  |
| Do the variances differ signif. ($P < 0.05$) | No | | | |
| ANOVA Table | SS | df | MS |  |
| Treatment (between columns) | 7442 | 2 | 3721 |  |
| Residual (within columns) | 6952 | 14 | 496.6 |  |
| Total | 14390 | 16 |  |  |
| Tukey's Multiple Comparison Test | Mean Diff. | q | P value | 95% CI of diff |
| Sham vs Vehicle 1ul | -5.8 | 0.582 | $P > 0.05$ | -42.69 to 31.09 |
| Sham vs Vehicle 2ul | -45.17 | 4.896 | $P < 0.01$ | -79.33 to -11.02 |
| Vehicle 1ul vs Vehicle 2ul | -39.37 | 4.267 | $P < 0.05$ | -73.53 to -5.216 |

B.
| Parameter | Value |  |  |
| --- | --- | --- | --- |
| Table Analyzed |  |  |  |
| left movement |  |  |  |
| Kruskal-Wallis test |  |  |  |
| P value | 0.0242 |  |  |
| Exact or approximate P value? | Gaussian Approximation |  |  |
| P value summary | * |  |  |
| Do the medians vary signif. ($P < 0.05$) | Yes | | |
| Number of groups | 4 |  |  |
| Kruskal-Wallis statistic | 9.422 |  |  |
| Dunn's Multiple Comparison Test | Difference | P value | Summary |
| Sham vs Vehicle 1ul | -2 | $P > 0.05$ | ns |
| Sham vs 6-OHDA 1ug | -4.6 | $P > 0.05$ | ns |
| Sham vs 6-OHDA 2ug | -11.2 | $P < 0.05$ | * |
| Vehicle 1ul vs 6-OHDA 1ug | -2.6 | $P > 0.05$ | ns |
| Vehicle 1ul vs 6-OHDA 2ug | -9.2 | $P > 0.05$ | ns |
| 6-OHDA 1ug vs 6-OHDA 2ug | -6.6 | $P > 0.05$ | ns |

C.
| Parameter | Value |  |  |  |
| --- | --- | --- | --- | --- |
| Table Analyzed |  |  |  |  |
| left movement |  |  |  |  |
| One-way analysis of variance |  |  |  |  |
| P value | 0.0657 |  |  |  |
| P value summary | ns |  |  |  |
| Are means signif. different? (P < 0.05) | No |  |  |  |
| Number of groups | 5 |  |  |  |
| F | 2.473 |  |  |  |
| R squared | 0.2479 |  |  |  |
| Bartlett's test for equal variances |  |  |  |  |
| Bartlett's statistic (corrected) | 9.039 |  |  |  |
| P value | 0.0601 |  |  |  |
| P value summary | ns |  |  |  |
| Do the variances differ signif. (P < 0.05) | No |  |  |  |
| ANOVA Table | SS | df | MS |  |
| Treatment (between columns) | 11610 | 4 | 2902 |  |
| Residual (within columns) | 35220 | 30 | 1174 |  |
| Total | 46830 | 34 |  |  |
| Tukey's Multiple Comparison Test | Mean Diff. | q | P value | 95% CI of diff |
| Sham vs Vehicle 1ul | -5.8 | 0.3785 | P > 0.05 | -68.65 to 57.05 |
| Sham vs 6-OHDA 2ug | -47.52 | 3.685 | P > 0.05 | -100.4 to 5.381 |
| Sham vs Vehicle+JB1a 3days | -18.31 | 1.291 | P > 0.05 | -76.50 to 39.88 |
| Sham vs 6-OHDA+JB1a 3days | -32.1 | 2.188 | P > 0.05 | -92.28 to 28.08 |
| Vehicle 1ul vs 6-OHDA 2ug | -41.72 | 3.235 | P > 0.05 | -94.61 to 11.18 |
| Vehicle 1ul vs Vehicle+JB1a 3days | -12.51 | 0.8822 | P > 0.05 | -70.70 to 45.68 |
| Vehicle 1ul vs 6-OHDA+JB1a 3days | -26.3 | 1.793 | P > 0.05 | -86.48 to 33.88 |
| 6-OHDA 2ug vs Vehicle+JB1a3days | 29.2 | 2.534 | P > 0.05 | -18.06 to 76.47 |
| 6-OHDA 2ug vs 6-OHDA+JB1a 3days | 15.42 | 1.273 | P > 0.05 | -34.27 to 65.11 |
| Vehicle+JB1a 3days vs 6-OHDA+JB1a 3days | -13.79 | 1.023 | P > 0.05 | -69.07 to 41.50 |

D.
| Parameter | Value |  |  |
| --- | --- | --- | --- |
| Table Analyzed |  |  |  |
| left movement |  |  |  |
| Kruskal-Wallis test |  |  |  |
| P value | 0.004 |  |  |
| Exact or approximate P value? | Gaussian Approximation |  |  |
| P value summary | ** |  |  |
| Do the medians vary signif. (P < 0.05) | Yes |  |  |
| Number of groups | 5 |  |  |
| Kruskal-Wallis statistic | 15.35 |  |  |
| Dunn's Multiple Comparison Test | Difference | P value | Summary |
| Sham vs Vehicle 1ul | -2.7 | P > 0.05 | ns |
| Sham vs 6-OHDA 2ug | -16.7 | P > 0.05 | ns |
| Sham vs Vehicle+JB1a 7days | -12.39 | P > 0.05 | ns |
| Sham vs 6-OHDA+JB1a 7days | -0.7556 | P > 0.05 | ns |
| Vehicle 1ul vs 6-OHDA 2ug | -14 | P > 0.05 | ns |
| Vehicle 1ul vs Vehicle+JB1a 7days | -9.688 | P > 0.05 | ns |
| Vehicle 1ul vs 6-OHDA+JB1a 7days | 1.944 | P > 0.05 | ns |
| 6-OHDA 2ug vs Vehicle+JB1a 7days | 4.313 | P > 0.05 | ns |
| 6-OHDA 2ug vs 6-OHDA+JB1a 7days | 15.94 | P < 0.05 | * |
| Vehicle+JB1a 7days vs 6-OHDA+JB1a 7days | 11.63 | P > 0.05 | ns |
| Table Analyzed |  |  |  |  |
| left movement |  |  |  |  |
| One-way analysis of variance |  |  |  |  |
| P value | 0.377 |  |  |  |
| P value summary | ns |  |  |  |
| Are means signif. different? (P < 0.05) | No |  |  |  |
| Number of groups | 4 |  |  |  |
| F | 1.094 |  |  |  |
| R squared | 0.1542 |  |  |  |
| ANOVA Table | SS | df | MS |  |
| Treatment (between columns) | 1745 | 3 | 581.7 |  |
| Residual (within columns) | 9569 | 18 | 531.6 |  |
| Total | 11310 | 21 |  |  |
| Tukey's Multiple Comparison Test | Mean Diff. | q | P value | 95% CI of diff |
| Sham vs Vehicle 1ul | -5.8 | 0.5625 | P > 0.05 | -47.01 to 35.41 |
| Sham vs Vehicle+6-OHDA | -22.23 | 2.391 | P > 0.05 | -59.37 to 14.92 |
| Sham vs JB1a+6-OHDA | -12.1 | 1.106 | P > 0.05 | -55.81 to 31.61 |
| Vehicle 1ul vs Vehicle+6-OHDA | -16.43 | 1.767 | P > 0.05 | -53.57 to 20.72 |
| Vehicle 1ul vs JB1a+6-OHDA | -6.3 | 0.576 | P > 0.05 | -50.01 to 37.41 |
| Vehicle+6-OHDA vs JB1a+6-OHDA | 10.13 | 1.014 | P > 0.05 | -29.78 to 50.03 |

**Supplementary Figure S4.**
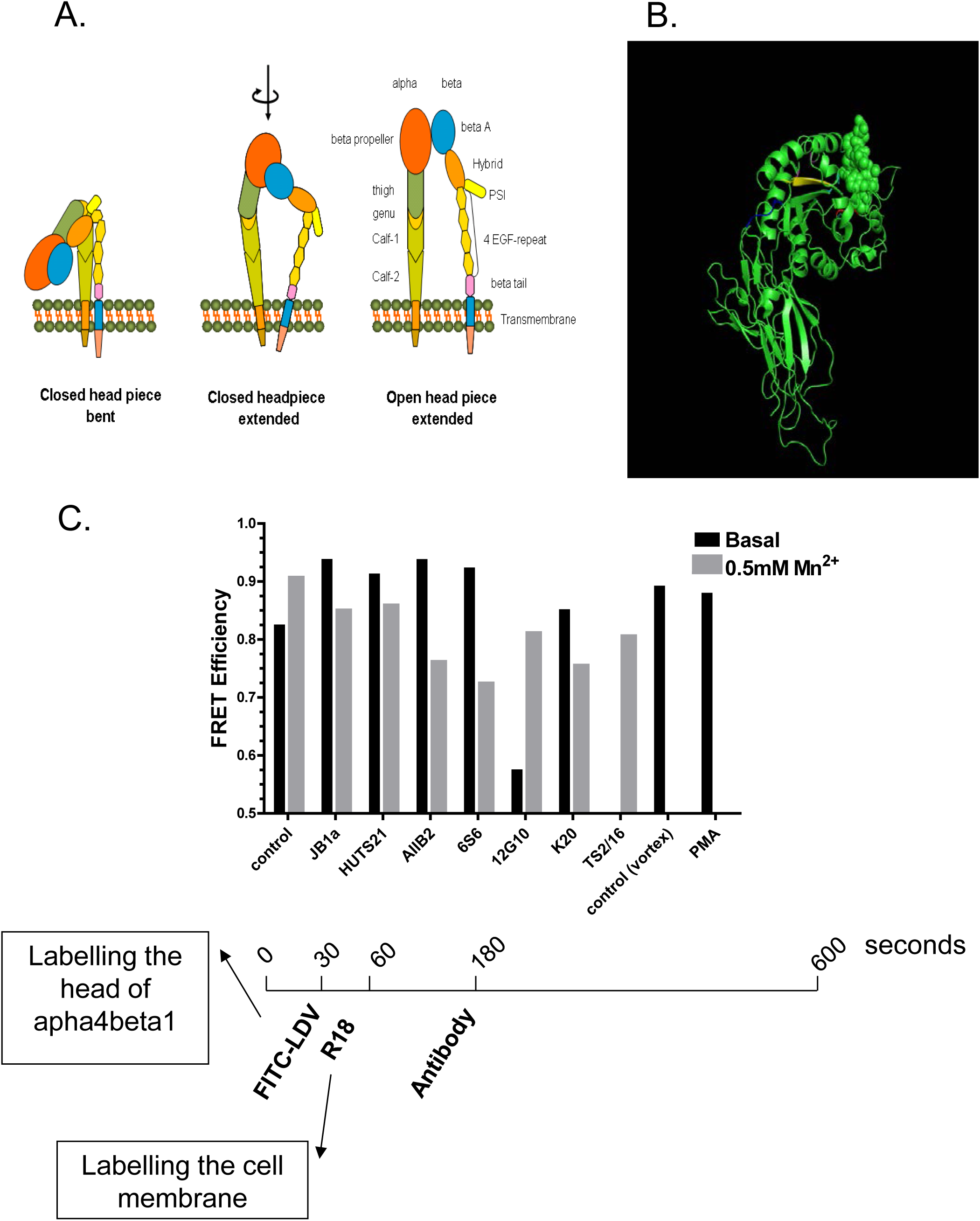
**A.** Schematic of β1 integrin conformational states (bent-closed, extended-closed, extended-open) (from Al-Jamal R, Harrison DJ, 2008). B. The JB1a binding site on the hybrid domain (residues 82–87), illustrating the proposed allosteric mechanism of action. C. Adapted FRET screening of the allosteric effect of β1 integrin antibodies. Methods as previously published by AlJamal-Naylor et al., 2012.

